# Complex genetic patterns in human arise from a simple range-expansion model over continental landmasses

**DOI:** 10.1101/158923

**Authors:** Ricardo Kanitz, Elsa G Guillot, Sylvain Antoniazza, Samuel Neuenschwander, Jérôme Goudet

## Abstract

Although it is generally accepted that geography is a major factor shaping human genetic differentiation, it is still disputed how much of this differentiation is a result of a simple process of isolation-by-distance, and if there are factors generating distinct clusters of genetic similarity. We address this question using a geographically explicit simulation framework coupled with an Approximate Bayesian Computation approach. Based on six simple summary statistics only, we estimated the most probable demographic parameters that shaped modern human evolution under an isolation by distance scenario, and found these were the following: an initial population in East Africa spread and grew from 4000 individuals to 5.7 million in about 132 000 years. Subsequent simulations with these estimates followed by cluster analyses produced results nearly identical to those obtained in real data. Thus, a simple diffusion model from East Africa explains a large portion of the genetic diversity patterns observed in modern humans. We argue that a model of isolation by distance along the continental landmasses might be the relevant null model to use when investigating selective effects in humans and probably many other species.

## Introduction

Departing from Africa around 100 kya (thousands year ago), modern humans colonized the globe, scattering over the continents. This slow migration process created genetic divergence as populations migrated, splitting along the way, to settle over the landmasses. The history of humans can be deciphered using genetic differences between populations, reaching further than anthropological knowledge [1]. With the increasing amount of genetic data, as well as the advance of theoretical models, historical and prehistorical processes playing a major role in shaping the observed genetic diversity can be better identified [2–4].

In particular it has been recognized that geography plays a major role in structuring populations [5]. The significance of geography as a driver of genetic diversity has already been demonstrated in many studies, for example in work based on blood group polymorphism [6], enzyme polymorphism [7], mitochondrial DNA complete sequences [8–10], and even complete genome sequences [11]. Acting as a barrier to migration, mountains and seas decrease the connectivity between populations, which correlates with genetic distance [3,12]. This monotonous relationship between (geographic) distance and diversity, known as cline, is expected under isolation by distance, in a continuous diffusion model. However, looking at populations worldwide, genetic patterns show clustering of populations into major groups (European, Asian, Melanesian, Native Americans and Africans) [12]. Although this continental split suggests the action of specific environmental or cultural forces, it remains unclear under which conditions these continental clusters emerge.

Hence, two types of patterns arise out of empirical population genetic studies, cline and cluster, which seems contradictory. Interpretations have flourished around these patterns, fueling the misplaced debate of human races [13,14].

Favoring a clinal view, some researchers have shown that human genetic variability declines as one moves further away from East Africa [4,15]. Moreover, it has been observed that there is a clear correlation (R^2^=0.85) between genetic distances (e.g., F_ST_) and geographic distances (along probable colonization routes). Although agreeing with this observed global pattern, studies favoring a cluster view point to discontinuities along the decline of diversity. For these clusters to appear, serial bottleneck events associated with isolation, must have generated what one could see as steps in a staircase of genetic diversity [3].

As an attempt to reconcile both perspectives, Serre et al. [2] brought the possibility that the geographically uneven sampling scheme used in most, if not all, worldwide studies on human genetics may have generated these clusters, which would merely reflect sampling bias. Rosenberg et al. [3] challenged this view taking advantage of an expanded dataset to argue that, among all other variables to be considered in the detection of clusters, geographic dispersion of samples has relatively little effect on the final outcome. In such cases, large amount of genetic data would always allow detecting discontinuities even if the distribution of sampled populations were completely uniform. Such discontinuities could be small, but still detectable and biologically relevant. Finally, another study, that focused on the geographical origin of modern humans, detected similar patterns of clines in F_ST_ and genetic diversity, and attributed the few deviations from these trends as being caused by “admixture or extreme isolation” [16]. Concretely, it remains unclear which underlying genetic and demographic processes could explain both cline and cluster observed pattern.

This apparent opposition between a clinal and a cluster view of human diversity arises because current models fails to re-create both patterns. Indeed those models tend simplify the complexity of human demographic history (population growths, migrations) as well as genetic processes (selection, drift). For example, studies looking for adaptation [17,18] as well as the association between genotype and phenotype [19] rely strongly on neutral models (diversity expected from drift and demography, no selection). Typically, some demographic scenarii create genetic polymorphisms which are indistinguishable from those supposedly left by selection. The deconvolution of selection and demographic signal is hindered by the lack of simple demographic model that would reproduce basic patterns of human diversity. For instance, Hofer et al. [20], looking at four continental human populations, detected an unexpected large proportion of loci (nearly a third of their database) with strong differences in allelic frequency. The authors suggested that the observed patterns are better explained by the combination of demographic and spatial bottlenecks with allele surfing in the front of range expansion rather than by selective factors [21]. In the allele surfing process, drift takes random samples of alleles at potentially different frequencies from the source population (i.e. founder effect), while the combination of range and demographic expansions amplifies this effect on the overall population by increasing the contribution of these alleles in the newly colonized regions. Therefore, to understand the recent genetic evolution of human populations, it is essential to have a good grasp on the demographic events underlying it. A first step to this end is to understand the spatial distribution of human genetic diversity and the emergence of strong discontinuities in empirical studies (i.e. formation of clusters).

To bridge the gaps between theoretical study and the discordance in empirical genetic studies, we present a simulation-based study. Here, we investigate the distribution of neutral genetic diversity in modern humans using spatially explicit simulations to model the demographic diffusion of our species throughout the globe and to recover the genetic signature left by this process. The simulations are used to estimate, the demo-genetic parameters best fitting a large microsatellite dataset of published data [22, 23] using Approximate Bayesian Computation (ABC) [24]. We do so by generating genetic data under a simple stepping stone model constrained by the shape of the continental masses. Based on the parameter estimates, we simulate a full dataset of individual genetic markers. We then compare simulated and empirical data using Principal Component Analysis (PCA) and analyses with the STRUCTURE software [25]. This permits to assess whether the proposed model is suitable for further population genetic studies, if it can generate patterns similar to the one observed in real data (clusters and cline). We then discuss the outcomes of such a model for the understanding of the processes defining human genetic diversity around the world and possible applications in the field.

## Material and Methods

### Empirical data

Data from this study represent a subset of the dataset originally made available by Pemberton et al. [23], Rosenberg et al. [3] and Wang et al. [22]. Since we used a strict mutation model, we chose 346 microsatellite loci whose length is proportional to the repeated segment length. These loci represent the ones termed ‘regular’ by Pemberton et al. [23] that are also available in the Wang et al [22] dataset. The number of populations in the original dataset was 78, totaling 1484 individuals distributed throughout the world (more details in Figure S1, Figure S2, and Table S1 in the Supplemental Data available online).

Although dense SNP datasets are available, we used a microsatellite dataset in this study for the following reasons: (i) The microsatellites used here have been extensively checked and shown to have equally sized repeat units, which is expected if they evolve under the stepwise mutation model; (ii) they are unlinked and essentially neutral; (iii) the number of samples and populations publicly available is greater than for SNPs [78 instead of 51 for the latter [26]] or whole genome (1000 genome project), and with better coverage of the American continent; and (iv) we could only simulate so many loci in a spatially-explicit approach with the currently available computational power. Note that being multiallelic markers, microsatellites contain more information per locus than SNPs [27].

### ABC

We estimated demographic and genetic parameters using an Approximate Bayesian Computation (ABC) framework. In brief, simulated dataset are generated over a large set of demographic parameters (start of expansion, initial population size, growth, etc.). The simulation outcome that best match the empirical data are selected to define a posterior probability distribution for each parameter. Genetic data were generated using a modified version of quantiNEMO [28] in a two-step process. First, individual-based forward-in-time simulations produce the demography of the expanding population. Then a backward in time coalescent-based process simulates the genetic polymorphism. Parameters were estimated using the ABC package ABCtoolbox [29].

For the demographic part, all simulations started at one single deme with a varying initial population size (N_i_, uniform prior distribution, from 2 to 5120), in Eastern Africa (9°1’48”N, 38°44’24”E) – today‘s Ethiopian city of Addis Ababa, the origin of human expansion as estimated by Ray et al. [30] and place of the oldest known modern humans remains [31]. The prior distribution for the time of the onset of this expansion had a normal distribution with mean of 155 000 years and standard deviation of 32,000 years (T, generation time of 25 years). These values were based on the combination of independently estimated dates of 141 455 ± 20 000 [32] and 171 500 ± 25 500 years ago [8]. These dates are more recent than the oldest reliably dated fossil remains in Ethiopia (195 000 ± 5000), which is expected since they most likely predate the spatial expansion of interest in this study [33]. Population regulation followed a stochastic logistic model [34] with intrinsic growth rate (r, lognormal prior, mean=0.5, SD=0.6) delimited by the deme‘s carrying capacity (N, uniform prior of 2-5120 individuals). Individuals are allowed to move between the four directly neighboring demes in a two-dimensional stepping-stone pattern with a given dispersal rate (m) sampled uniformly between 0 and 0. 5. Genetic data were generated using a coalescent approach to simulate genealogies for 20 microsatellite loci (single stepwise mutation model) with a mutation rate μ (uniform prior of 10^−5^-10^−3^ mutations/locus/generation) for the same 70 populations and same number of individuals as the observed sampling scheme (see Table S2).

### Summary statistics

In ABC, summary statistics are used to compare observations with simulations [24,35]. Ideally, these summaries should be a set of a small number of measures that maximize the information. Initially, we explored a large set of different summary statistics: number of alleles, allelic richness [36], Garza-Williamson‘s M [37] and gene diversity [38] per sampled population; pairwise F_ST_ [39] and Chord-distances [40] between samples. Considering that many of them did not bring extra information to our inference scheme, while hindering the estimation [41], we used two different techniques to reduce the dimensionality of the dataset. We retained a subset made of the 2,415 pairwise F_ST_ between populations and the number of alleles (A) for each of the 70 demes. These 2,485 summary statistics were then transformed into six “pattern” statistics, summarizing the relationships between F_ST_, number of alleles and geographic distance as follows: The number of alleles sample was regressed on the geographic distance between the sampled location and Addis Abeba, and pairwise F_ST_ were regressed against pairwise geographic distances. From these two regressions, we extracted six pattern statistics, namely the means, slopes, and the logarithm of the sum of residuals. The calculations of summary and pattern statistics for the observed data were carried out in R and the R-package *hierfstat* [42]. Finally, these six pattern statistics were used for the estimates of the demo-genetic parameters and subsequent validations. We also used partial least squares (PLS) to reduce the original 2,485 summary statistics to a small number of components [43]. This technique gave very similar (but no better) results for the validations and a few parameters had slightly different estimated values (Figure S4). In the main text, we only report the results obtained with the six pattern statistics.

### Estimates

The six simulations parameters (N_i_, μ, m, N, r, T) were estimated based on a comparison of the simulated and the observed summary statistics and a subsequent estimation step. The comparison of the summary statistics was obtained by assessing the Euclidean distance between simulations and the statistics from the empirical data, which can be used to rank the simulations from closest to most distant from the observations. Here, we retained the 5,000 simulations with smallest Euclidean distances from the observations. This subset of simulations was then used to estimate the parameter values using a weighted generalized linear model (GLM) [44] of the six pattern statistics with the ABCtoolbox software [29].

### Validation

In order to assess the quality of our estimation process, we perform a standard ABC validation. Hence, we used pseudo-observed values taken from the simulations. We quantify how well these values could be recovered when estimated through our ABC pipeline [45]. This was done for 1000 different pseudo-observations for each of the six investigated parameters. We calculated then the correlation (R^2^) for the regression between pseudo-observed and estimated values, the slope of this regression, the standardized root mean squared error of the mode (SRMSE) and the proportion of estimates for which the 95% higher posterior density interval included the true value.

### Full-dataset simulations

Using these estimated parameters we generate new simulated samples with 100 loci per individual, with quantiNEMO. To investigate the effect of our estimated parameters, we ran three sets of 100 simulations each whose parameter values were sampled from the (i) prior distribution of the estimation step, (ii) posterior distribution (95%HPD) of the estimation step or (iii) taken directly from the point estimates (mode values of the posteriors) of the estimation step. Using the output of these simulations, we investigated how well these simulations could reproduce analyses carried out on the real data set. To check for consistency, the first comparison was based on the same six pattern statistics used for the estimations (i.e. mean, slope and sum of residuals for number of alleles and pairwise F_ST_). A second comparison was based on the first two axes of a principal component analysis (PCA) computed on the individual allele frequencies in each sampled population. Since the sign of the coordinates along PCA components can differ between replicates, we compared the different sets of simulations by means of the squared correlation between observed and simulated PCA results. Each axis was considered separately. Thus, for each simulation, we estimated an R^2^ representing the correlation between simulated and observed populations coordinates on the PCA axes. These R^2^ values were compared across the three different sets of simulations (Prior, 95%HPD and Mode).

Finally, we ran a clustering analysis using STRUCTURE v2.3.4 [25] on the point estimate simulated set. Each simulation was analyzed for varying K (the number of clusters) between 1 and 7. Each structure analysis was run for 250 000 iterations, discarding the first 50 000 as burn-in. To assess the accuracy of our model, we ran STRUCTURE on the empirical data, but for these analyses we used the whole set of 346 microsatellite loci and ran 25 replicates for each K. We processed the STRUCTURE outputs with CLUMPP [46] in order to align the different replicates to compare the simulations data with the observations. We also carried out the estimation of the number of groups (K) best explaining the variation present in simulations and observations following Evanno et al. [47]. The ΔK was estimated based on 25 replicates for each STRUCTURE run.

## Results

We ran in total 1,183,831 simulations based on prior distributions; 974,934 (82.4%) successfully colonized all the sampled patches and were therefore used in the subsequent analyses. We obtained posterior estimates for all six demo-genetic parameters, which are presented in Table 1 (point estimates; for their complete distributions, see Figure S3). The inferred distribution of each parameter presents a clear unique peak, as expected under a good estimation. Briefly, we estimate a first expansion 132 kya with an initial population size close to 4,000 individuals, expanding with a growth rate of 0.149 and a migration rate of 0.041. The mutation rate μ is estimated at 2.6x10^−4^ mutation/site/generations.

**Table 1:** Accuracy table and estimates of the six variable parameters inferred by the ABC framework. Point estimate corresponds to the mode of the posterior distribution, while HPD95% interval represents the parameter values comprised within the 95% higher posterior density interval. R^2^ stands for the coefficient of determination of pseudo-observed on estimated values; SRMSE is the root mean squared error of the mode, standardized between 0 and 1; Prop. HPD95% stands for the proportion of tests for which 95% higher posterior density intervals include the true value. All rates are per generation (25years).

| | T (years) | $N_i$<br>(ind.) | N (ind.) | $\mu$ | r | m |
| --- | --- | --- | --- | --- | --- | --- |
| Point estimate | 132 250 | 3952 | 5 725 656 | $2.6 \times 10^{-4}$ | 0.149 | 0.041 |
| HPD95% interval | 60 850 -<br>203 900 | 920 -<br>5120 | 35 658 - 20<br>905 776 | $9.3 \times 10^{-5}$ -<br>$4.4 \times 10^{-4}$ | 0.036 -<br>0.679 | 0 - 0.177 |
| $R^2$ | 0.235 | 0.399 | 0.431 | 0.877 | 0.286 | 0.57 |
| SRMSE | 0.132 | 0.233 | 0.227 | 0.099 | 0.108 | 0.187 |
| Slope | 0.248 | 0.536 | 0.602 | 0.908 | 0.352 | 0.682 |
| Prop.<br>HPD95% | 0.993 | 0.956 | 0.981 | 0.977 | 0.983 | 0.979 |

To assess the accuracy of these inferred parameters, we used a validation procedure [29] based on 1000 independent simulations. The mutation rate (µ) estimation is satisfactory since we observed a strong correlation between pseudo-observations and estimations (R^2^=0.877) for which the slope was nearly 1 (slope=0.908), and the error rate low (SRMSE=0.099). The proportion of the estimates that included the pseudo-observed value within their 95%HPD interval was 0.977, suggesting that our posteriors are slightly conservative. Good inference was also achieved for migration rate (m), current population size (N) and initial population size (N_i_) for which the R^2^ values were about 0.5 and the slopes above 0.6. We had rather poor estimations for time of the onset (T) and population growth rate (r) where R^2^ values were below 0.3 (Table 1).

Full-dataset simulations. The posterior estimates above were then used in further simulations to create three sets of simulated genetic markers (100 simulated microsatellite loci), mimicking the empirical sampling scheme. These additional simulations were carried-out by randomly sampling parameter values from either (i) the prior posterior, (ii) the truncated posterior (at the 95%HPD level) distributions or (iii) the point estimates. As these parameters were estimated using basic genetic polymorphism summary statistics, it is essential to check whether such simple expansion can produce the empirical cline and clustering patterns.

We first verified that our simulations were able to replicate the clinal pattern observed in the original genetic data. Figure 1A shows the empirical cline with a reduction of genetic diversity while increasing geographic distance, while Figure 1B shows in comparison the simulated cline using point estimates parameters. In both cases, the general pattern is the same: a steady reduction of diversity for populations as one moves away from Addis Ababa, and a clear-cut increase of genetic differentiation with geographic distance.

**Fig 1:**
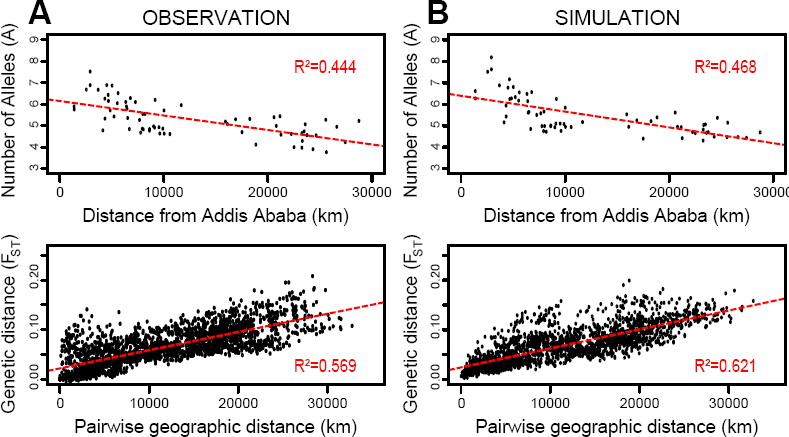
Comparison of the patterns of isolation by distance generated with the empirical and simulated data. In A, the patterns obtained for the observed data; in B, the result of one of the simulations based on the point estimates. Each point represents a population (top) or a pairwise population comparison (bottom); the dashed lines represent the linear regressions of these points (whose R^2^ values are informed).

The comparison of the three simulation sets and the empirical cline emphasizes the power of the ABC inference. Indeed, as expected, parameters sampled from posterior distribution produce patterns closer to the empirical dataset than the prior distribution. The points estimates produce patterns close, on average, to the posterior distribution, with less variation around the true value (Figure 2 and Figure S5). Finally the cline produced by the set of point estimate simulations is very close to the empirical cline.

**Fig 2:**
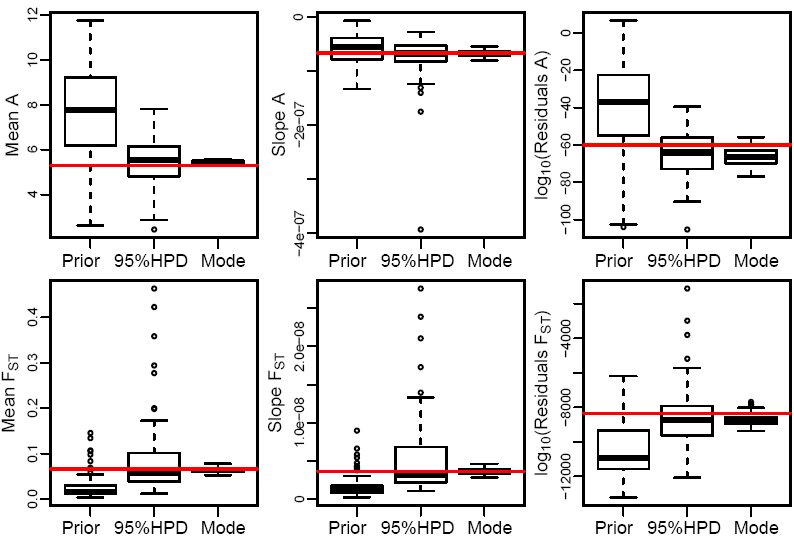
Distribution of estimated statistics from three simulated dataset and empirical observation (horizontal gray line). Within each plot, we present the different sources for the simulations that generated the distributions: “Prior” are simulations sampled randomly from the whole prior; “95%HPD” are simulations run based on the 95% higher posterior density estimates for all parameters; and “Mode” represent simulations based on the point estimates for all parameters.

Next, we investigated whether the simulated genetic data could reproduce the clustering patterns observed in the Principal Component Analysis (PCA). In the empirical dataset, one observes clear divisions between continental groups (Figure 3A), as previously demonstrated elsewhere [9,48]. The PCA results based on our simulations returned a pattern very similar to that observed (Figure 3A). The convergence of the estimation of parameters, from prior, to 95% HPD, to point estimates, can also be assessed looking at the PCA. The correlation between observation and simulations in their principal components (PC1 and PC2) are presented in Figure 3B. The data simulated under the 3 scenarios generated patterns for the first PCA component extremely similar to what is observed in the real data set. For the second PCA component, the similarity to the observed pattern was small for dataset generated under the prior parameters distribution, and increased for data simulated with the posterior parameters distribution and point estimates.

**Fig 3:**
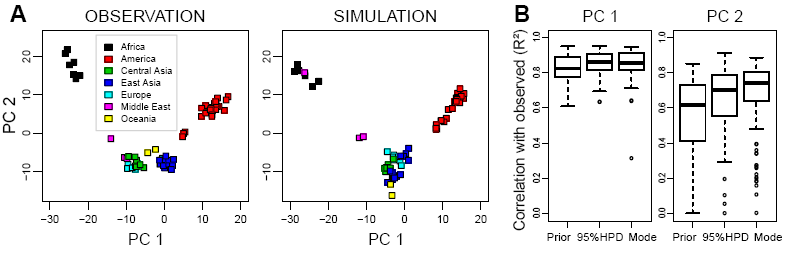
PCA results in real observation and simulations. A, Comparison of PCA applied to the empirical data (left) and one selected simulation (right). The first (PC 1) and second (PC 2) principal components are represented here, where each point represents one of the analyzed populations, grouped by continents. B, Boxplots of the correlation values between the two first principal components in observations and simulations based on the prior distribution (“Prior”), 95% higher posterior density distribution (“95%HPD”), and on the point estimates (“Mode”).

Finally, we also looked at the partitioning pattern generated by the software STRUCTURE. Simulations and empirical data gave the same estimates of the most likely number of groups (K) within the worldwide sample either using the highest likelihood of the data as the criteria for defining K (which led to K=7 in both observations and simulations), or using ΔK [47], which favored K=2 both for observations and simulations (Figure S7). The similarities also persist in the way the different individual genomes are allocated to the different clusters resulting from this analysis. They generated, for both empirical and simulated data, remarkably similar results for K=2 to K=4 (Figure 4). For K=2, we observe a cluster of Africans and a cluster of Americans whereas all other individuals are admixed of these groups to different extent; the proportion of admixture obtained for the different individuals in the simulations matches almost perfectly with that seen in the observation. For K=3, Eurasian populations emerge from the previous African cluster with a few differences between simulations and observation: In the observations, Middle-Easterners and Europeans group with Africans; whereas in the simulations, they are admixed between the African and East Asian clusters. For K=4, the sub-Saharan samples split from the rest of the world creating a cluster unique to Africans. While for the empirical observation this division is very clear, the results based on the simulated data show a more gradual pattern with Middle-Eastern and European mixed-ancestry samples. Beyond K=4, the patterns observed between simulations and observations diverge: while single populations start to emerge as separate clusters in the observation; higher values of K lead to the appearance of admixed individuals and populations within the already existing groups, creating no new clusters (Figure S6). Interestingly, in both simulations and observation, the grouping pattern is relatively consistent with the continental partitioning of the populations.

**Fig 4:**
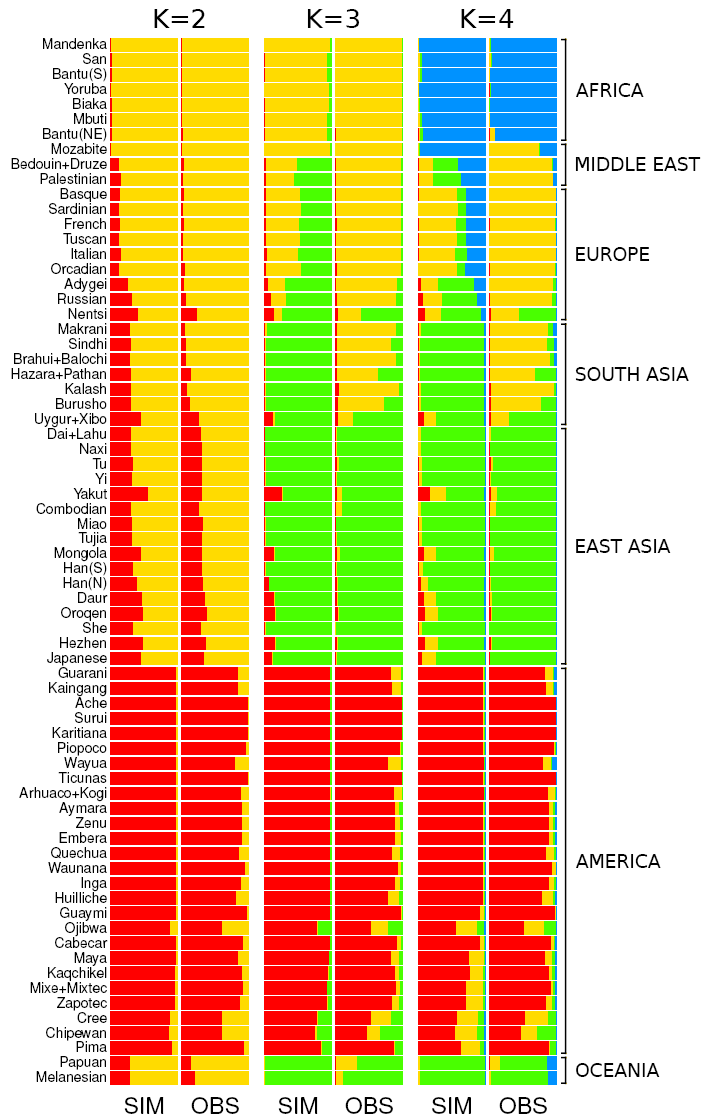
Comparison between the STRUCTURE results obtained for observed (OBS) and simulated (SIM) data. Horizontal bars represent the 70 populations as used in the simulations and the different shades of gray code for the proportion of each inferred ancestry group (K from 2 to 4).

## Discussion

We have shown using approximate Bayesian computation that a simple model of expansion from East Africa using the world-wide landmasses leads to meaningful estimates of the past demography of our species. Furthermore, when genetic data sets generated according to this past demography are analyzed with Principal component analyses or the Structure program, we obtain results that are extremely similar to those observed in the original human microsatellite dataset. We discuss these findings below

Despite the increasing use of genetic markers in anthropological reconstruction, it remains unclear how to model the observed patterns of genetic diversity around the world, largely because of the complexity of evolutionary processes of the human species. Specifically, the apparent opposition between cline and clustering patterns, as observed in empirical studies, remains a challenge as most existing model fail to reproduce both patterns. Owing to the release of new fast simulation tools, such as quantiNemo, and the rising availability of global datasets, we reconstruct a simple expansion scenario that reproduces the clustering effect of modern populations, using large samples of published microsatellites data.

Based on empirical data of 346 microsatellites in 1,484 individuals from 70 populations, this study has inferred six parameters (T, N_i_, N, μ, r, m) that defines a worldwide expansion model using the computationally intensive ABC framework. Despite the simplicity of the model, the inference works remarkably well. The estimated values are similar to other studies. The mutation rate (µ=2.6 10^−4^ mut/allele/gen) matches recent estimates [49]. The growth rate (r=0.149) is close to rates described elsewhere when applying logistic growth to humans [30]. We inferred a start of expansion from Addis Ababa around 132 kya, close to previous estimates [32]. Moreover, the validation, based on the estimation of known parameters using simulated pseudo-observation, confirms the accuracy of the inferred values.

The inferred demic expansion model along landmasses generates genetic patterns very similar to those observed in the real dataset. Similarly, to other studies [9], these simulations confirm the signatures of isolation-by-distance and constant decrease of genetic diversity with increasing distances from Addis Ababa. Strikingly, these similarities are robust towards the inferred parameters, as tested with three simulation sets (parameters issued from prior distribution, posterior distribution or point estimates).

To investigate clustering patterns, PCA and STRUCTURE analyses were performed. The PCA on the simulated dataset shows a strong correlation with both the first and second principal components calculated from the observation. The STRUCTURE analysis presents closely related results between real data and simulations: the number of groups which better explains the diversity in the samples is the same for both. The population division for up to four clusters remains very similar. Hence, this study shows the possibility to reproduce both observed isolation by distance and continental clusters under a unifying model of simple expansion.

To understand the underlying processes reproducing this pattern, it is interesting to have a close look at the partitioning analyses. PCA has long been used in human population genetics [50], it relates genetic variation to the geographic distribution of populations [51] and individuals [52]. Simulated and empirical data are similarly scattered on the two first principal components. The coordinates of the samples along the first axis (Figure 3B) show a very high correlation with the observed coordinates, even for simulations based on the prior, uninformative, distribution of the parameters. This indicates that the first component of the PCA (capturing the largest fraction of the genetic variance) probably relates to the origin of the expansion (which occurs in the same place, East Africa, for all simulations) and demic diffusion. The second principal component seems to be more sensitive to the choice of the parameter values, the correlation between observation and simulations increasing when the parameters used for the simulations get closer to the estimation.

Although admixture-based analyses are not completely independent from PCA [53], the most surprising result obtained in this study comes from the population clustering analysis in STRUCTURE. Indeed, no previous study has shown the appearance of clusters from a simple diffusion process such as that we used in our simulations. In fact, based on ΔK, the estimation of the best number of groups, allowing for admixt individuals, is consistent between simulated and empirical data with K=2 which suggests weak support to separate genetic groups. In both cases, the assignment of each populations to the clusters is extremely similar, the model is therefore able to reproduce the overall genetic patterns.

However, global population genetic studies have been – regardless of the previous finding – consistently analyzed as if continental clusters were relevant [3]. Hence, we overlook the lack of significance of multiple partitioning on worldwide samples to analyze the data with K>2; the apparition of continental clustering is investigated in the simulations. The American populations are the first to stand out; second, a separation between European and African versus East Asian; and then the Africans alone stand out from the rest. There are a few exceptions though. The Mozabite population, from North Africa, tends to group with the other African populations in the PCA results for the simulations; while, in the observed data, they group with the Middle-Eastern and European populations. It is possible that more recent events of contact through the Strait of Gibraltar [54] or the Fertile Crescent, which are not captured by our simulations, contributed to this discrepancy. Another explanation could be the absence of the potentially important barrier of the Sahara Desert in the simulations, which may have played an important role in isolating North Africans from sub-Saharan populations. Although previous studies have modeled such environmental heterogeneity [30] it is extremely difficult to model environmental changes, like the expansion of Sahara, through the last 100,000 years. Moreover, the simulated European/Middle-Eastern populations are admixed unlike the empirical data, which may be caused by the absence of the Sahara as well. Other studies have shown that the peopling of Europe, the Fertile Crescent and North Africa is more complex than a simple expansion [1,55]. Despite these few (albeit important) discrepancies, this very basic model reproduces the global worldwide patterns remarkably well.

A potential bias in this study appears with the use of microsatellite loci which have a higher polymorphism than the more popular SNP data which are becoming standard. However, unlike SNP that are affected by ascertainment bias, evolutionary models of microsatellite data are better known. Moreover, the amount information captured with a limited number of loci, constraining the speed of simulations, is higher in microsatellites. Hence to grasp any bias introduced by the type of markers we provide a comparison of previous studies across these two kinds of markers. For the PCA results, studies on SNP worldwide datasets [48,52,56] return results very similar those obtained here both for the empirical and simulated data (Figure 3A). The first component correlates with the distance from the start of expansion, with Americas being the furthest. The second axis correlates with a north south geographical separation. For the STRUCTURE analyses, the clustering pattern remains similar across markers. Indeed, Rosenberg et al. [3] using STRUCTURE on microsatellite data have found results very similar to those obtained with SNPs in Li et al. [9], which are, in turn, very similar to our results in Figure 4. Therefore, for capturing the overall human genetic distribution, the SNP data may increase the resolution of the results, but does not seem to affect the general patterns that are replicated in the model we propose here.

The results obtained here shed new light on the “cline vs. clusters” controversy. The fact that a simple model of two-dimensional dispersion on a homogeneous world succeeds in producing results so similar to the real data in many different analyses is strong support for an overall clinal view of the distribution of human genetic diversity over the globe. Even though the simulations used here involve some sophistication, the underlying model is simple and can easily be considered in further population genetics studies: isolation-by-distance and continuous decline of diversity as we move away from East Africa. These two patterns are easily described by two linear regressions after all.

The clinal model for the global distribution of human diversity encounters support in other biological and cultural systems. Skull morphological diversity, for example, shows a clear and steady decline within population diversity as the distance from Africa increases and is in perfect agreement with what is found in DNA [57]. Language, a cultural feature, also shows a similar pattern. Distance from Africa, alone, explains 30% of the reduction in phonemic diversity as measured in 504 languages worldwide [58].

This view of human genetic diversity distributed over a continuous cline reinforces the inadequacy of biological “races” as clear separation between different human groups [14,59–61]. Nonetheless, classifying humans in different groups is still common practice in many genetic studies (mostly medical genetics) [62]. Typically, cluster identification can be useful for localized studies. At a smaller scale, a refinement of the genetic specificity of populations can be linked to the local prevalence of some conditions such as the identification of some genes involved in malaria resistance or lactase persistence [63]. However, if one is interested in broader scale studies, separating individuals by continents may lead to mistakes and misconceptions, as each individual genetic marker will be better defined by cline.

Working against the current trend of always more intricate models that capture a maximum of variation in the data, but failing to reproduce the global genetic patterns of cline and clusters, we present here a very simple expansion scheme over continental landmasses. Although additional spatial heterogeneity could help to improve this basic neutral model (e.g. by accounting for the Sahara), the simple one used here proved to be very useful for explaining the main patterns of human genetic variation. Such a model may represent a good choice for establishing a neutral background in future studies looking at more complex questions in modern human evolution such as the detection of selective events [64–66]. Specifically, its simplicity permits large scale fast simulations necessary for quantitative analysis of genetic markers. Indeed, the more specific models of local individual movement are not able to produce the vast amount of simulations needed for statistical analysis [67]. Moreover, with added complexity comes a vast set of added parameters (for example, local migration, time of demographic events, spatial heterogeneity). Although these may seem more biologically significant, these models tend to over-fit the data, as the information contained in the genetic markers may not be sufficient to infer a large set of parameters. A bigger number of inferred parameters also decreases the power of ABC while increasing exponentially the computation time. The good fit of this very simple model over the dataset argues for using expansion-diffusion models or more simply isolation by distance, instead of discrete populations, as a fundamental model of human population genetics.

## Acknowledgements

We would like to thank Nelson J. R. Fagundes for commenting on previous versions of this manuscript and John Pannell for fruitful discussions on the subject. Also, we are grateful to Anna Kostikova who independently verified the calculation of geographic distances between populations.

The computations were performed on clusters provided by Vital-IT (http://www.vital-it.ch) Center for high-performance computing of the Swiss Institute of Bioinformatics (SIB).

